# Resident memory T cell development is associated with AP-1 transcription factor upregulation across anatomical niches

**DOI:** 10.1101/2023.09.29.560006

**Authors:** Neal P. Smith, Yu Yan, Youdong Pan, Jason B. Williams, Kasidet Manakongtreecheep, Shishir Pant, Jingxia Zhao, Tian Tian, Timothy Pan, Claire Stingley, Kevin Wu, Jiang Zhang, Alexander L. Kley, Peter K. Sorger, Alexandra-Chloé Villani, Thomas S. Kupper

**Author notes:** Alexandra-Chloé Villani, Thomas S. Kupper.

## Abstract

Tissue-resident memory T (T_RM_) cells play a central role in immune responses to pathogens across all barrier tissues after infection. However, the underlying mechanisms that drive T_RM_ differentiation and priming for their recall effector function remains unclear. In this study, we leveraged both newly generated and publicly available single-cell RNA-sequencing (scRNAseq) data generated across 10 developmental time points to define features of CD8 T_RM_ across both skin and small-intestine intraepithelial lymphocytes (siIEL). We employed linear modeling to capture temporally-associated gene programs that increase their expression levels in T cell subsets transitioning from an effector to a memory T cell state. In addition to capturing tissue-specific gene programs, we defined a consensus T_RM_ signature of 60 genes across skin and siIEL that can effectively distinguish T_RM_ from circulating T cell populations, providing a more specific T_RM_ signature than what was previously generated by comparing bulk T_RM_ to naïve or non-tissue resident memory populations. This updated T_RM_ signature included the AP-1 transcription factor family members *Fos, Fosb* and *Fosl2*. Moreover, ATACseq analysis detected an enrichment of AP-1-specific motifs at open chromatin sites in mature T_RM_. *CyCIF* tissue imaging detected nuclear co-localization of AP-1 members *Fosb* and *Junb* in resting CD8 T_RM_ >100 days post-infection. Taken together, these results reveal a critical role of AP-1 transcription factor members in T_RM_ biology and suggests a novel mechanism for rapid reactivation of resting T_RM_ in tissue upon antigen encounter.

## Introduction

Adaptive immune memory mediated by T cells is central to host defense, and our appreciation of its complexity has evolved considerably over the last half century. Unanticipated heterogeneity in cytokine production by memory T cells was introduced in 1986, and migratory heterogeneity in circulating memory T cells was introduced in 1999 with the description of human central and effector memory T cells (T_CM_, T_EM_)^1,2^. Over the past 15 years, an additional population of resident memory T cells (T_RM_) has been the focus of many studies. T_RM_ cells have been shown to reside long-term in peripheral tissues, rather than circulate through blood or secondary lymphoid organs, and play a critical role in antigen-specific recall immune responses^3^. They also appear to play a role in antagonizing tumor growth^4,5^. Studies in mice have suggested that T_RM_ populations arise from early effector T cells, which after entry and conditioning in peripheral tissues alter their phenotype to maintain residency and acquire long-lasting memory^6^. Recently this view has been challenged by the idea that some effector T cells are already committed to a T_RM_ lineage before they enter tissue ^7^. Our ability to discriminate between these possibilities and our understanding of the gene program changes that lead to these unique functions remains incomplete.

To date, regulation of a series of key transcription factors has been associated with T_RM_ development and establishment. It has been well described that Hobit or its human analog Blimp1 are key regulators of T_RM_ maintenance that function via repression of genes associated with tissue egress^8^. *Runx3* has been associated with T_RM_ establishment specifically in CD8 T_RM_, and is responsible for a cell’s responsiveness to the TGFβ signals associated with T_RM_ differentiation ^9,10^. Conversely, T_RM_ precursors have been shown to lose expression of transcription factors that regulate tissue egress and lymph-homing molecules, including *Tcf7, Eomes* and *Klf2* ^6,11^. More recently, an elegant study utilizing single-cell RNAseq (scRNAseq) by Kurd *et al.* described how the expression of AP-1 transcription factor members *Junb* and *Fosl2* as well as *Nr4a2* are essential for the development of T_RM_ in the small intestine^12^. The authors hypothesized that these AP-1 family members down-regulate the expression of T-bet, given that a similar mechanism exists in T helper (Th17) cells ^13^.

CD8 T_RM_ development and biology have been studied extensively in a number of infectious mouse models that employ transgenic T cells responsive to viral epitopes such as Vaccinia virus (VACV) and Herpes simplex virus (HSV) in skin as well as influenza in the lung ^8,14–21^. Much of our additional knowledge on CD8 T_RM_ in different tissues has emerged from similar mouse studies using lymphocytic choriomeningitis virus (LCMV) infection^10,12,22^. Unlike VACV, HSV and influenza, LCMV infection generates a systemic rather than a local immune response, leading to the formation of T_RM_ across many tissues, including gut, liver, lung, kidney, and salivary glands ^23–28^. While the local responses created by viruses such as VACV and the systemic responses induced by LCMV represent distinct biological phenomena, comparisons of the T cell differentiation processes in these two complimentary systems can help us better define traits that are common to tissue and circulating T cell development.

Teleologically, CD8 T_RM_ that are induced through VACV skin infection are generated to protect against future re-infection through skin. In this setting, both antigen-driven activation and tissue-specific trafficking education are initiated in the skin draining lymph node (dLN) microenvironment, the first site of antigen contact with naïve T cells. While we have previously reported bulk transcriptional profiles of activated cells in such dLN^17^ this profiling strategy does not capture biological heterogeneity defining the T cells derived from the dLN in the early time points post-infection. To overcome these limitations, the present study leveraged scRNAseq with OT-I T cells from the skin and dLN across a time course spanning 0-60 days post-infection to study in more detail the evolution of T_RM_ and T_CM_ in parallel from the same infection. The granularity and cellular resolution provided by such scRNAseq strategy performed across a time course enabled us to define previously uncharacterized heterogeneity among cells from the dLN in the early timepoints post-infection. Additionally, our experimental approach utilizing a time course allowed for downstream linear modeling. This analysis captured gene programs associated with temporal T_RM_ development and defined genes associated with T_RM_ and T_CM_ cell fates in both our VACV-induced skin T cells, as well as publicly available LCMV-induced T cells in the small intestine. We identified AP-1 transcription factor family members as key genes contributing to our consensus T_RM_ signature across tissue compartments, which were confirmed to be associated with mature resting T_RM_ cells via ATACseq and highly multiplexed, tissue-based CyCIF microscopy^29^. In particular, the high expression and nuclear localization of AP-1 members in resting T_RM_ is unique among memory T cell lineages. These findings provide new insights into the different trajectories of T cells after antigen exposure.

## RESULTS

### scRNAseq of OT-I cells reveals 13 T cell subsets paving the way for memory T cell development across time and tissue source

To investigate the process of antigen-specific T cell development, OT-I transgenic mouse T cells were adoptively transferred into recipient mice one day before skin infection with a recombinant VACV that expresses chicken ovalbumin peptide (amino acids 257–264) under the control of an early gene promoter (rVACV-OVA) (**Methods**). Activated OT-I effector T cells were readily found in the skin as early as 5 days after infection and reached their maximum level at day 10 (*data not shown*), before beginning to decrease in number, as previously reported ^21^. scRNAseq was performed on FACS-sorted OT-I cells from both skin and dLN, respectively, at serial timepoints from days 0-60 (**Figure 1A, Extended data** **Figure 1A; Methods**). After filtering out contaminating populations that lacked expression of canonical T cell markers (*Cd3d, Cd8a, Trac*), we recovered 76,028 high-quality cells across dLN and skin (**Extended data** **Figure 1B**). Surprisingly, cells from the skin across all timepoints were divided into two main populations, the major one of which that was defined canonical CD8 T cell markers (termed “skin1”; *Cd3d, Cd8A, Trbc2*) and a minor one that had additional expression of MHC-II related molecules (*Cd74, H2-Aa*), Ig receptors (*Fcer1g, Fcgr2a*) and *Tyrobp* (termed “skin2”; **Extended** **figure 1C-E**). Flow cytometry of day 30 OT-I cells from an independent experiment confirmed a population of T cells with skin2 markers (**Extended data** **figure 1F**). Given concerns about the skin2 population representing T-cell:mononuclear phagocyte (MNP) doublets, a commonly observed scRNAseq pitfall^30^, Imagestream flow cytometry was employed to validate the existence of this skin2 subset. The results were informative; while there was indeed a population of doublets - where T-cell markers were expressed on cells distinct from those expressing Mhc-II, Tyrobp and CD74 - a population of single OT-I cells that expressed skin2 markers was also clearly observed. (**Extended data** **Figure 1G**). Given this observation, we hypothesized that our skin2 population represented a mixture of T cell/MNP doublets as well as a distinct population of T cells derived from antigen-activated naïve T cells. However, given our inability to separate out a pure population of skin2 T cells from the doublets, this small skin2 population as well as other contaminating doublets (n = 12,763 cells) was removed from downstream analysis for the present paper (**Extended data** **Figure 1H**), resulting in a filtered dataset of 63,265 cells that were re-clustered (**Methods**).

**Figure 1.**
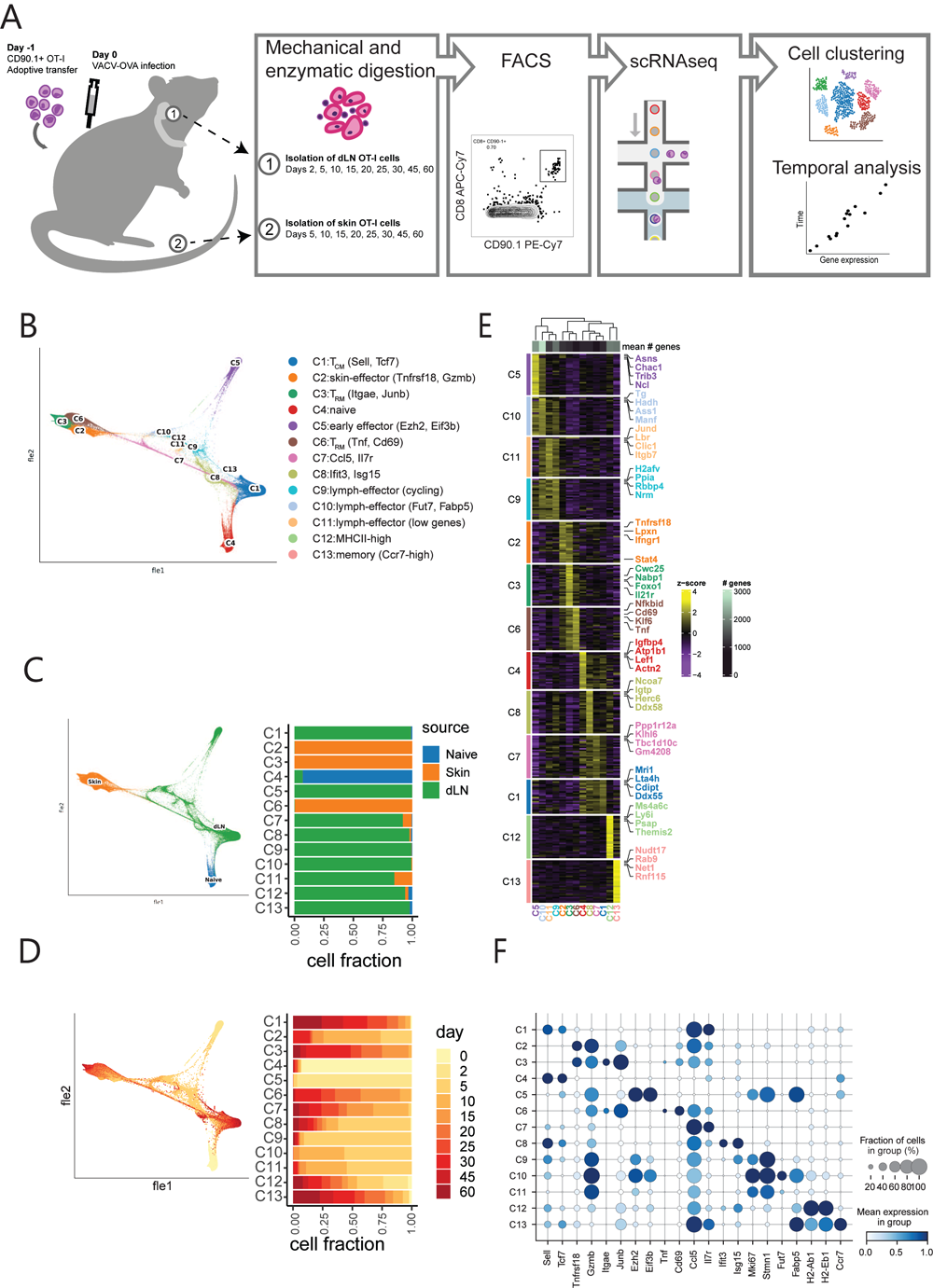
**(A)** Schematic of the experimental design. **(B)** Force-directed layout embedding (FLE) of 63,265 high-quality single cells, colored by predicted Leiden cluster listed on the right. **(C)** (left) FLE embedding of cells pseudocolored by tissue source. (right) Source composition of every cluster. Bars represent the fraction of cells in every cluster that were derived from the corresponding source. **(D)** (left) FLE embedding of cells pseudocolored by experimental timepoint. (right) Timepoint composition of every cluster. Bars represent the fraction of cells in every cluster that were derived from the corresponding timepoint. **(E)** Heatmap showing the top discriminative gene sets for each cell cluster, compared to every other cluster. Color scales denote the normalized gene expression (mean zero, unit variance) for each cluster and the mean number of genes captured per cluster (top bar). **(F)** Dot plot showing the percentage (size of the dot) and scaled expression (color) of known T-cell subset marker genes.

When visualized in low-dimension space, our filtered dataset showed uniform expression of *Cd3d, Cd8a* and *Trbc2,* with no contaminating non-T cell populations (**Extended data** **Figure 2A; Methods**). To compare the data across both tissue sources and time-point, we performed unbiased clustering analysis which resulted in 13 distinct cell populations (**Extended data** **Figure 2B**). The defined subsets were distinguished by different anatomical sites and timepoints, reflecting the emergence of different T cell subsets over time (**Figure 1B-D, Extended data** **Figure 2C, D**). To best visualize the trajectory of our cells, we used Force-directed layout embedding (FLE), an approach recommended for temporal data^31^. This dimensionality reduction approach grouped cells in order of timepoints measured – capturing the differentiation of naïve T cells to effector and memory cells – while also maintaining distinctions created by tissue sources and Leiden clustering (**Figure 1B-D**). All subsets were defined by a distinct set of genes using statistically complementary strategies (area under the curve [AUC] ≥ 0.75, one-vs-all [OVA] pseudobulk false discovery rate [FDR] < 0.05; see **Methods**), justifying our cluster resolution (**Figure 1E; Supplementary table 1**). C4 was almost exclusively comprised of naïve T cells and showed expression of lymph-homing markers *Sell* and *Ccr7* (**Figure 1F**). We identified a subset of T_CM_ cells (C1) that shared many phenotypic markers with naïve cells but came from late timepoints (**Figure 1D**), likely reflecting the similar quiescent state of naïve and T_CM_ cells. There was an additional subset of dLN memory cells with high *Ccr7* expression (C13), but they represented a very small proportion of the captured cells (133 cells, 0.21%) (**Extended data** **figure 2E**).

**Figure 2.**
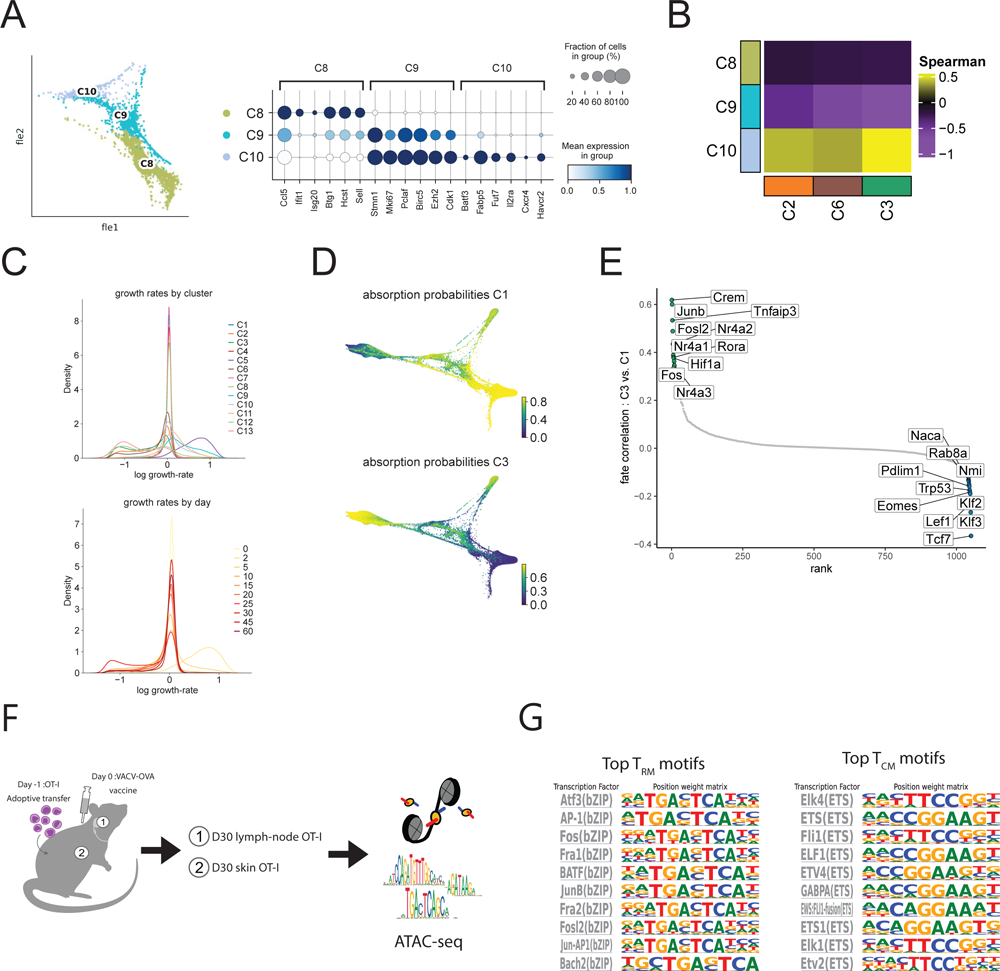
**(A)** (left) FLE of clusters most associated with day 5 dLN cells (C8, C9, and C10). (right) Dot plot showing the percentage (size of the dot) and scaled expression (color) of select marker genes for each of the 3 clusters. **(B)** Pairwise Spearman correlation between the OVA log2 fold-change values of clusters C8, C9 and C10 versus C2, C6 and C3. **(C)** A growth rate was calculated by comparing the relative expression of genes involved in proliferation versus apoptosis. Histograms show distribution of this growth rate across clusters. **(D)** Absorption probabilities for cells to be a part of the C1 terminal macrostate (left) or C3 terminal macrostate (right). Color scale represents degree of fate priming. **(E)** Transcription factors most associated with each Waddington-OT determined terminal cell state were defined. The top 10 transcription factors associated with C3 terminal state (left, top) and C1 terminal state (right, bottom) are labeled. **(F)** Schematic of the ATACseq experimental design. ATACseq schematic inspired by Bentsen *et al*.^78^ **(G)** HOMER-known motif analysis comparing T_RM_ and T_CM_ samples profiled. Shown are the transcription factors and position weight matrices for the top 10 known motifs for T_RM_ (left) and T_CM_ (right).

Five dLN populations (**Figure 1C, Extended data** **Figure 2C**) were identified as having an effector-like phenotype that broadly expressed markers of cytotoxicity (e.g. *Gzma, Gzmb*) (**Figure 1F**). These populations included C5 early effector cells that were largely made up by day 2 cells and were defined by markers known to be associated with early antigen-activation such as *Ezh2* and *Eif3b*^32,33^. Additional populations of dLN effector T cells included the C8 population expressing high levels of interferon stimulated gene (ISG) signature (e.g. *Ifit3*, *Isg15*), the C9 population defined by high levels of cell cycle markers, the C10 population marked by the expression of the known skin-homing marker *Fut7* and T_RM_ -associated gene *Fabp5* and the C11 population that seemed to be defined by having a lower number of genes detected, possibly reflecting lower-quality cells (**Figures 1E-F**, **Extended data** **figure 2F**). Over 75% of cells from C8-11 populations were from day 10 or earlier (**Figure 1D**). A small population of dLN cells defined by high expression of MHC-II machinery (C12) was also present in the data.

Unlike the cells with a similar expression profile in skin that were filtered out, these represented a small fraction of the total data (n = 449 cells, 0.71%) (**Extended data** **figure 2E**). Given their small representation and our inability to distinguish doublets, this population was not explored further.

In the skin, three distinct populations were defined (**Figure 1C, Extended data** **Figure 1C**). C2 represented the earliest of the skin T cells with >50% of cells from days 5 and 10 (**Figure 1D, Extended data** **Figure 2E**), with an effector T cell phenotype defined by *Tnfrsf18, Gzmb* and *Ctla4* expression **(Figure 1E; Supplementary table 1)**. In contrast, C3 and C6 were defined as T_RM_ cells because they were made up largely from the later timepoints (> day 15) and nearly absent in the earliest timepoints (≤ day 10). C3 expressed the highest level of *Itgae* and *Icos*, while C6 expressed higher levels of *Cd69* and *Tnf*. Interestingly, C3 and C6 were represented in nearly equal proportions in the middle timepoints (days 15-25: C3 = 43.7% of skin cells, C6 = 44.0% of skin cells) (**Extended data** **figure 2E**). However, C3 became the dominant T_RM_ population by the end of the time course (days 45-60: C3 = 87.2% of skin cells, C6 = 12.2% of skin cells), suggesting that C3 represented the terminal resting T_RM_ population.

### T cell subsets with skin-homing features defined in lymph-node

We sought to assess in which tissue compartment (dLN versus skin) and at which time point does T cell subset differentiation start emerging through the course of T cell development. Our first two timepoints revealed little heterogeneity, with 96% of Naïve day 0 cells being found in C4 and 98% of day 2 cells belonging to C5 (**Extended data** **Figure 2E**). Interestingly cell subset diversification was first observed at day 5 across three distinct clusters (C8, C9 and C10) representing 85% of day 5 cells from the dLN (**Extended data** **Figure 2E, 3A**). C8 was enriched in interferon-response genes (*Ifit1, Isg20*) as well as *Btg1*, a gene associated with T cell quiescence^34^ (**Figure 2A, Extended data** **Figure 3B**). Additionally, C8 had the highest expression of the lymph-homing molecule *Sell*. Subsets C9 and C10 were enriched for cell cycle markers (*Stmn1, Mki67, Pclaf, Birc5*), while only C10 expressed genes associated with skin-homing and T_RM_ development (*Fabp5, Fut7*). Interestingly, the presence of C8 and C9 cells persist through our time course (C8: 21% >D15 cells, C9: 3.7% >D15 cells), while C10 is almost completely absent by day 15 (0.1% > D15 cells), suggesting C10 is a transient cell state in the dLN (**Extended data** **Figure 2E**). To understand the relationship between these dLN populations (C8, C9 and C10) and the defined skin cell subsets (C2, C6 and C3), we used each cluster’s gene expression profile to calculate pairwise Spearman correlations (**Figure 2B; Methods**), which showed that only the expression profile of C10 correlated with C2, C6 and C3. These results support the hypothesis that the emergence of cell subset diversification among antigen-activated T cells appears prior to trafficking to peripheral tissues.

**Figure 3.**
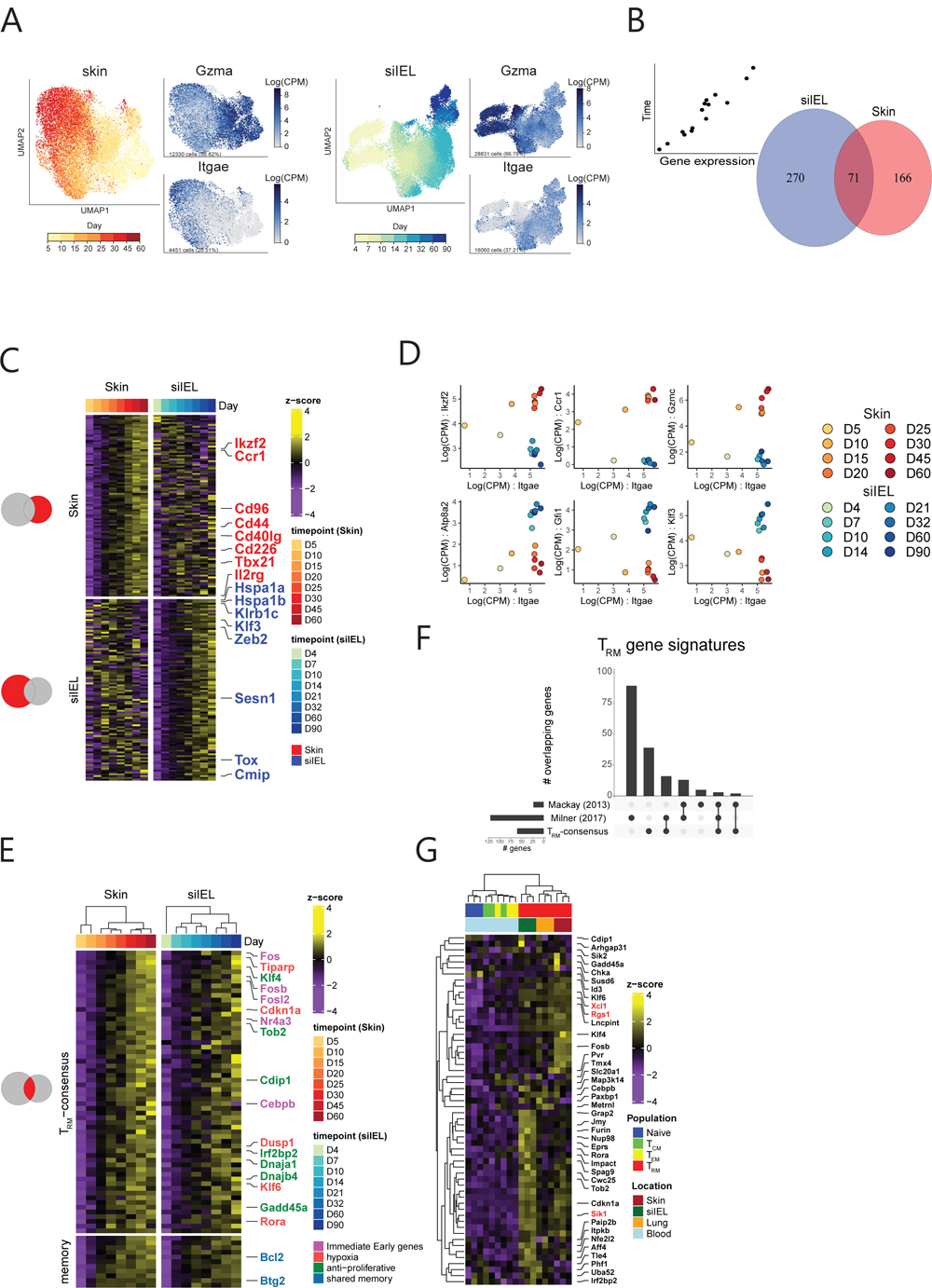
**(A)** Uniform Manifold Approximation and Projection (UMAP) embedding of skin (left) and publicly available siIEL scRNAseq data (right) pseudocolored by experimental timepoint. To the right of each timepoint UMAP are feature plots using color to indicate gene expression levels (Log(CPM)) of *Gzma* and *Itgae*. Cell number and percentage represent expression across cells of all timepoints within the given anatomical niche. **(B)** Venn diagram of the significant T_RM_ -associated genes in siIEL (left) and skin (right) as determined by linear modelling. **(C)** Heatmap showing the top 100 genes unique to the skin (top) and siIEL (bottom) T_RM_ signatures. Top bar indicates the associated timepoints. Color scales denote the normalized gene expression (mean zero, unit variance) for each timepoint. **(D)** Scatter plots showing the Log(CPM) of select skin-specific (top) and siIEL-specific (bottom) T_RM_ genes on the y-axis and Log(CPM) of *Itgae* on the x-axis. Color scale indicates both anatomical location and timepoint the sample was from. **(E)** Heatmap showing the consensus T_RM_ gene signature across timepoints in both skin (left) and siIEL (right) datasets. Color scale denotes normalized gene expression (mean zero, unit variance) for each timepoint. The genes on top represent those unique to the T_RM_ signature genes (n = 60), while the genes on the bottom represent those additionally found in the T_CM_ signature (n = 11). **(F)** UpSet plot showing the overlap between our consensus T_RM_ 60-gene signature and two previously published signatures by Milner *et al.* (2017) and Mackay *et al.* (2013). Each column represents a unique intersection as shown by the dark points in the dot-matrix. Bars for each column represent the size of the overlap between each combination. Bars on left represent the size of each unique T_RM_ gene set. **(G)** Heatmap showing expression levels of our consensus gene signature across T-cell subset samples publicly available from Mackay *et al.* 2013^20^. Expression levels were derived from Microarray data. Color scales denote the normalized gene expression (mean zero, unit variance) for each sample. Genes listed in black are those unique to our consensus T_RM_ signature. Genes listed in red are those that are shared among all 3 T_RM_ signatures.

### AP-1-family transcription factors are associated with T_RM_ development

To better understand the development of terminally differentiated T cell fates, we employed the Waddington-Optimal Transport (Waddington-OT) algorithm through the CellRank suite of tools^35,36^ (**Methods**). Waddington-OT infers temporal couplings between cells profiled across our experimental time-course, capturing the transcriptional programs and regulators driving the transition between cell states ^35^. To determine if the predicted rate of cellular proliferation should be considered when modeling cellular trajectories, we calculated a growth rate based on expression of genes associated with the cell cycle and apoptosis (**Figure 2C**). The predicted growth rates across the 13 clusters and 10 timepoints were fairly uniform with the exception of C5, which was largely made up of day 2 cells and most associated a higher growth rate (mean log growth rate of C5: 0.64, mean log growth rate of all other clusters: -0.13). However, given that this cluster represented only small a fraction of our dataset (7,130 cells, 11.3%), we opted for modeling the data with a uniform growth rate.

When using the Waddington-OT algorithm to predict two terminal macrostates (regions of the phenotypic manifold that cells are unlikely to leave), the subsets identified as the most likely to represent terminally differentiated cells were C1 and C3, which correspond to late-time point dLN (T_CM_) and skin cells (T_RM_), respectively (**figure 2D**). To elucidate drivers of terminal differentiation, we looked at the transcription factors most associated with either the C1 or C3 lineages (**Figure 2E; Supplementary table 2; Methods**). While the C1 trajectory was associated with transcription factors known to maintain T cell quiescence (*Klf3*), lymph homing, and T cell memory (*Tcf7, Eomes*) ^37,38^, the C3 trajectory strongly correlated with AP-1 transcription factor members *Junb*, *Fosl2*, and *Fos*, along with other additional known immediate early genes *Nr4a1*, *Nr4a2*, and *Nr4a3*.

To investigate further the transcriptional regulators driving memory T cell differentiation, we performed Assay for Transposase-Accessible Chromatin-sequencing (ATACseq) on D30 post VACV skin infection T cells, including skin T_RM_ and dLN T_CM_ cells (**Figure 2F**, **Extended data** **Figure 1A, 3C; Methods**). We probed for differences in transcription factor motifs between these two T cell subsets using HOMER motif analysis and consistent with our transcriptional signatures, we found the T_RM_ cells to be strongly enriched with bZIP family transcription factor motifs including AP-1, *Fos, JunB* and *Fosl2*, validating the regulators predicted by the Waddington-OT algorithm. Conversely, T_CM_ were enriched for ETS family transcription factor motifs, including those corresponding to *Ets1, Elk1 and Elk4* (**Figure 2G; Supplementary table 3**).

### Gene programs leading to resident-memory state in Skin and siIEL T cells include distinct features

To better understand genes that are associated with T_RM_ differentiation, skin T cells were subclustered independently for further downstream analysis. Additionally, we included in our analysis a previously published dataset that used similar time course kinetics to analyze the differentiation of siIEL T_RM_ cells using a LCMV infection model^12^. We leveraged these two temporal datasets to better define the distinct and shared features of T_RM_ differentiation across distinct anatomical niches ^12^ (**Figure 3A**). To identify genes associated with T_RM_ development, a linear model was fit to gene expression data, capturing genes that gradually increase from the early timepoints to later time points in each tissue (**Methods**). This approach defined 341 and 237 T_RM_ genes in the siIEL and skin respectively (FDR < 0.1, regression slope > 0.2; **Figure 3B, Supplementary table 4**). The majority of T_RM_ genes were unique to a single anatomical site (siIEL = 270 unique genes, skin = 166 unique genes). For example, the transcription factor *Ikzf2* (Helios; *p*-value = 5e-4) was specific to skin but not siIEL (**Figure 3C-D**). There were additional immune mediators specific to skin, including *Ccr1* (*p*-value = 0.001) and *Gzmc* (*p*-value = 0.0001). In the siIEL compartment, there was an enrichment of heat-shock proteins (HSPs) associated with T_RM_ development (*Hspa1a*, *Hspa1b, Hsph1, Hsp90aa1, Dnaja4*) which have not been described in this context. Of note, HSPs play a critical role in the adaptation to hypoxic conditions, which have recently been associated with T_RM_ differentiation^39^. Other siIEL T_RM_ defining transcripts included *Atp8a2*, (*p*-value = 0.002) a gene recently reported to be associated with siIEL regulation as well as transcription factors *Klf3* (*p*-value = 2e-6), *Tox* (*p*-value = 0.0001) and *Gfi1* (*p*-value = 0.003) ^40^. Gene set enrichment analysis (GSEA) was then performed using both the HALLMARK and KEGG databases to identify gene sets that are associated with T_RM_ development. Skin T_RM_ development was distinctively associated with apoptosis and IL2 signaling pathways while gene sets associated with siIEL T_RM_ development included those for Wnt and Notch signaling (**Extended data** **Figure 4A-B; Supplementary table 5**). This analysis also revealed 13 shared T_RM_ gene sets, including the Hallmark Hypoxia (skin NES = 1.90, *p*-value = 0.0001; siIEL NES = 1.47, *p*-value = 0.007) and TGF beta signaling (skin NES = 2.19, *p*-value = 0.0001; siIEL NES = 1.80, *p*-value = 0.001) pathways (**Extended data** **Figure 4C, Supplementary table 5**), suggesting shared core programs leading to resident-memory states in skin and siIEL T cells.

**Figure 4.**
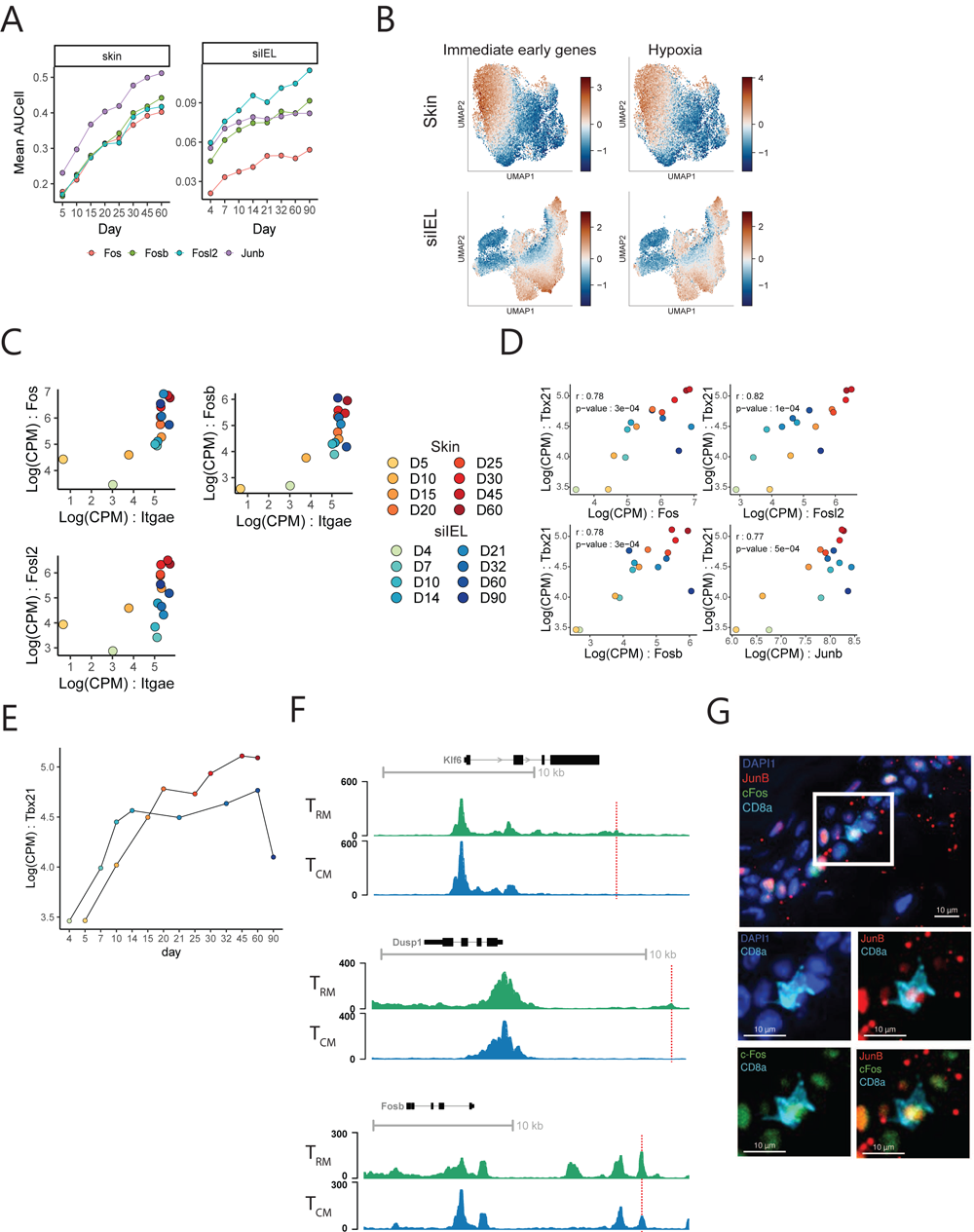
**(A)** Mean SCENIC Aucell scores for select regulons over time in the skin (left) and siIEL (right). Each line represents a unique regulon and the points represent the mean AUCell score for the regulon at the experimental timepoint. **(B)** Gene set scoring of skin (top) and siIEL (bottom) T cells with an immediate early (left) or hypoxia (right) gene set. **(C)** Expression of Fos-family genes versus *Itgae* over time in skin and siIEL. Each point represents a sample detailed in the legend that is shared with **(D)** and **(E)**, and the X-and Y-axes represent the Log(CPM) of *Itgae* and Fos family members respectively. **(D)** scatter plots showing the Log(CPM) of *Tbx21* on the y-axis and Log(CPM) of *Fosl2, Fos, Fosb,* and *Junb* on the x-axis across skin and siIEL timepoints. Color scale indicates both anatomical location and experimental timepoint from which the sample came from. r and p-values are from Pearson correlation. **(E)** *Tbx21* expression in skin and siIEL over time. The X-axis represents time while the Y-axis represents Log(CPM) of *Tbx21*. Dots are connected by their neighboring timepoints. **(F)** ATACseq tracks from our T_RM_ and T_CM_ samples at the *Klf6, Dusp1* and *Fosb* loci. Dotted line represents location of predicted Fos binding motif enriched in T_RM_ versus T_CM_. **(G)** t-CyCIF image of mouse tail skin epidermis 154 days after rVACV-OVA vaccination. Duplex and composite images of highlighted CD8+ T_RM_ cell expressing CD8 (turquoise), JunB (red), and cFos (green).

### dLN and spleen T cells from VACV and LCMV models share transcriptional programs that lead to circulating memory T cell development

To better understand the development of circulating memory T cells (T_CIRC_) across different viral infection systems, we employed the same linear modeling approach as described above with dLN cells from our VACV infection time-course and from the publicly available time course dataset looking at the differentiation of T_CIRC_ in the spleen after LCMV infection^12^ (**Extended data Figure 5A-B; Methods**). This approach defined 270 and 773 dLN-and spleen-T_CIRC_ genes respectively (FDR < 0.1, regression slope > 0.2; **Supplementary table 4**). Unlike the distinct gene profiles observed between skin and siIEL T_RM_ cells, the majority of dLN T_CIRC_ genes from the VACV model were shared with spleen T_CIRC_ genes from the LCMV model (n = 208 genes, 77%) (**Extended data Figure 5B**). Notably, the majority the dLN-specific genes in the spleen time-course were most highly expressed at the latest timepoints (**Extended data figure 5C)**. This would suggest that, while strict statistical criteria did not allow them to be considered T_CIRC_ genes in the spleen, many may still be involved in T_CIRC_ differentiation in dLN and spleen. In contrast, several T_CIRC_-specific genes found in the spleen of LCMV-infected mice were not up-regulated in the dLN T_CIRC_ of VACV-infected mice (n = 565 genes) (**Extended data Figure 5B**). This could plausibly be attributed to distinct cellular compositions of the T_CIRC_ populations at each anatomical site. While the dLN will predominantly contain T_CM_ cells, the spleen comprises a mix of T_CM_ and T_EM_.

### Skin and siIEL T cells from VACV and LCMV models share core transcriptional programs essential to resident memory T cell development

One of our goals in modeling T_RM_ development across multiple anatomical compartments was to curate a consensus T_RM_ gene signature. While many of the defined T_RM_ genes were distinct in our two anatomical niches, we found 71 genes were commonly expressed among both skin and siIEL compartments (**Figure 3B, 3E; Supplementary table 4**; FDR < 0.1, regression slope > 0.2). Of these 71 genes, 60 genes were unique to T_RM_ and 11 genes were also included in our T_CIRC_ signature. We predict these genes are important for general T cell memory and include *Btg2* and *Bcl2,* factors known to mediate T cell quiescence and T cell memory ^34,41^. Previous studies generated gene sets associated with T_RM_ development by comparing fully differentiated T_RM_ to other differentiated memory T cell subsets (T_CM_, T_EM_). To highlight the strength of our temporal linear modeling to derive a T_RM_-related gene signature, we compared our results to two previously published T_RM_ gene signatures^10,20^ (**Figure 3F**). Of the 60 genes found in our consensus T_RM_ signature, 21 were found in at least one of the other gene sets, with 3 genes being found across all gene signatures (*Xcl1, Sik1, Rgs1*). 39 genes were unique to our approach, including *Id3,* a transcription factor that has been associated with T cell memory ^42^ and the immediate early genes *Fosb* and *Cebpb*. Importantly, hierarchical clustering of naïve, T_CM_ and T_EM_ microarray samples^20^ based on the 60 gene consensus T_RM_ signature perfectly segregated these populations from T_RM_, further validating the use of linear modeling to capture genes defining T_RM_ along a temporal trajectory (**Figure 3G**). Noteworthy, when we examine the genes unique to the largest current T_RM_ gene set^10^ (n = 121 genes) that was generated with microarray data, their expression patterns do not consistently track with T_RM_ development in the skin and siIEL (**Extended data Figure 6A**). In addition to the different analytical approaches taken to generate our consensus T_RM_ gene set versus the previously published gene sets, many discrepancies may be due to the differences in technologies used to examine gene expression, as certain genes are not well captured by scRNAseq.

Previous work has demonstrated that the regulation of key transcription factors are important for T_RM_ development, such as *Runx3, Notch1* and *Zfp683*^8–10,43^. To define T-_RM_-associated transcription factors that are both tissue-specific and shared across the skin and siIEL compartments, a modified version of the SCENIC algorithm was used to analyze the skin and siIEL scRNAseq datasets (**Extended data Figure 6B-C; Supplementary table 6;** methods) ^44,45^. Briefly, SCENIC was used to calculate AUCell scores for specific regulons, which are defined as a transcription factor and potential targets of that transcription factor. These scores were then used as input to a linear model analogous to the approach used to capture gene expression patterns in **Figures 3B-E**. The skin-specific regulons identified (FDR < 0.1) included *Klf6* (*p*-value = 8.5e-5) *Irf4* (*p*-value = 0.004), *Nfil3* (*p*-value = 1.2e-5) and *Spi1* (*p*-value = 0.001). Several siIEL-specific regulons were also identified (FDR < 0.1) that included *Foxo3* (*p*-value = 0.001), *Elf2* (*p*-value = 0.01), and *Myc* (*p*-value = 0.04), which have all been reported to either inhibit proliferation and/or have been associated with T_RM_ development in an LCMV _model12,46,47._

### AP-1 transcription factor family members define T_RM_ across different anatomical niches

Interestingly, the SCENIC analysis identified 12 regulons common to both skin and siIEL T_RM_ development, including the AP-1 transcription factor members *Fos* (skin *p*-value = 1.5e-5; siIEL *p*-value = 7e-4)*, Fosb* (skin *p*-value = 3e-6; siIEL *p*-value = 2e-4)*, Fosl2* (skin *p*-value = 1.4e-5; siIEL *p*-value = 1e-4) and *Junb* (skin *p*-value = 3e-5; siIEL *p*-value = 0.008) (**Figure 4A, Extended data Figure 6C; Supplementary table 6**).When considering the functional relevance of our 60-gene consensus T_RM_ signature, we noticed an enrichment of genes associated with hypoxia signaling (*Rora, Cdkn1a, Klf6, Dusp1*), a known environmental factor that can induce a T_RM_ phenotype^39^, as well as genes associated with Immediate early responses (*Fos, Fosb, Fosl2, Nr4a3, Cebpb*), which include AP-1 transcription factor family members (**Figure 3E, 4B-C, Extended data Figure 7A**). A prevailing hypothesis about the role of AP-1 transcription factor members in T_RM_ development is that they coordinate the down-regulation of the transcription factor T-bet (encoded by the gene *Tbx21*) ^12,48^. Surprisingly, however, *Tbx21* itself met our stringent criteria for a T_RM_-specific gene in skin (regression slope = 0.32, *p*-value = 4.39e-10) and almost met our criteria in siIEL (regression slope = 0.15, *p*-value = 0.002) (**Figure 3C**). Furthermore, expression of *Tbx21* showed a strong positive correlation with the AP-1 family members *Fos* (r = 0.78 *p*-value = 3e-4), *Fosl2* (r = 0.82, *p*-value 1e-4), *Fosb* (r = 0.78, *p*-value = 3e-4) and *Junb* (r = 0.77, *p*-value = 5e-04) across both skin and siIEL, suggesting the proposed negative regulation of T-bet by AP-1 members to enable T_RM_ development is unlikely (**Figure 4D-E**). Nonetheless, our ATAC-seq analysis identified open-chromatin sites with predicted *Fos/Junb* binding motifs unique to T_RM_ around the loci of specific T_RM_ consensus signature genes, which included *Klf6, Dusp1 and Fosb* (**Figure 4F**).

Given the expression of AP-1 family members in mature T_RM_, coupled with an enrichment of motifs for bZIP family members as determined by ATAC-seq analysis, we endeavored to examine T_RM_ in tissue by CyCIF^29^ (**Methods**). Specifically, we used high-plex tissue imaging to determine whether AP-1 family member proteins could be identified in skin OT-I T_RM_ cells >100 days post-infection and whether they were localized to the cytoplasm or the nucleus (**Figure 4G**). While CD8 was clearly diffusely expressed consistent with membrane localization, JunB and c-Fos staining were co-expressed and found in the nucleus (co-localized with the nuclear Hoechst stain). To date, this is the first demonstration of expression of JunB and c-Fos in the nucleus of resting memory T cells. This suggests that AP-1 complexes are preformed and poised to be activated in T_RM_.

## Discussion

In this study, we sought to carefully examine the ontogeny and development of different memory T cells arising from a common naïve T cell population by performing a scRNAseq time course analysis of CD8 OT-I T cells after a skin infection with VACV. Our analysis revealed that cellular diversification among antigen-activated CD8 T cells arises early in the dLN, with the emergence of a subset of T cells expressing lymph-homing or skin-homing molecules, respectively. Through the use of linear modeling to predict genes associated with T_RM_ development over time, we derived a consensus T_RM_ signature shared by skin and siIEL that was marked by an overrepresentation of AP-1 transcription factor family members. We provide several orthogonal lines of evidence supporting the role of AP-1 members in T_RM_ development through our Waddington-OT trajectory analysis as well as follow-up ATAC-seq and microscopy results that validated these AP-1 targets. Building on these results, future work will be needed to decipher the mechanism of action of AP-1 family members in the context of T_RM_ recall immune responses.

Our initial observation of two skin T-cell populations – one (termed “skin1”) defined by conventional markers of T-cells (*Cd3d, Cd8a, Trbc2*) and another (“termed skin2”) defined by additional up-regulation of antigen-presentation machinery (*Cd74, H2-Aa*) and Ig-receptors (*Fcer1g, Fcgr2b*) – was an unexpected finding. It is conceivable that the “skin2” population captures T cell:MNP doublets; the sequencing of doublets is commonly observed in droplet-based scRNAseq experiments and can be challenging to interpret without the use of orthogonal approaches to validate findings^49^. We therefore used imaging flow cytometry to further investigate these “skin2” cells and validated the existence of single cells co-expressing CD8, MHC-II and CD74 among day 30 T_RM_ cells. A subset of Natural-Killer cell-like circulating T cells expressing *FCER1G* and *TYROBP* (similarly expressed by “skin2”) have been described in humans and this population was found to have anti-tumor activity, but to our knowledge, such a population has not been described in peripheral tissues or animal models ^50^. More recently, Pallett *et al.* described T cells acquiring unexpected cell surface markers from antigen-presenting cells through trogocytosis, a possibility that we cannot completely rule out when considering our imaging results ^51^. As expected, imaging flow cytometry also confirmed the presence of T cell:MNP doublets, which were captured among the events with the highest expression of CD74. Currently, given that we were unable to separate these doublets from the biologically distinct group of singlets in our single-cell data, together with our inability to rule out trogocytosis as a putative explanation for the imaging results, we were prompted to remove the entire “skin2” subset of cells from downstream analysis. However, future work will need to address the identity and function of the non-doublet “skin2” T cells further validated via imaging flow cytometry to assess whether these cells represent a biologically distinct T_RM_ subset. Looked at another way, it is revealing that a potentially novel T cell population may be hidden in a cluster of doublet cells; this should serve as a cautionary tale for analyzing scRNAseq data. Notably, one should be careful in interpreting doublet clusters having genes shared by multiple cellular lineages without further pursuing orthogonal validation like Amnis flow. Doing this may enable the disentangling of truly novel biology from doublet technical artifacts found in scRNAseq technologies.

Our analysis of cells isolated from the dLN showed that T cell subsets derived from naïve cells showed cellular diversification as early as day 5. Intriguingly one of the three dLN subsets (C10) found predominantly at day 5 exclusively expressed *Fut7*. This gene has been shown to be necessary for T-cell trafficking to the skin by glycosylating ligands for E and P selectin in the context of tumor-clearance, leading us to speculate that this population represents the cells destined to traffic to the skin ^52^. Additionally, the gene expression profile of C10 most closely correlated with skin clusters C2, C3 and C6, supporting the hypothesis that these are skin-trafficking cells. In contrast, among the early dLN populations, C8 showed the highest expression of *Sell* as well as *Btg1*, a gene recently associated with T cell quiescence ^34^. These results support the hypothesis that activation in the dLN leads to the generation of different T cell subsets, some of which appear to be destined to traffic to the skin, while others are destined to remain in the lymph node or recirculate between blood and dLN.

Our comparative analysis of transcriptional programs driving T_RM_ ontogeny across anatomical sites found 270 and 166 genes that were unique to siIEL and skin compartments, respectively. The siIEL-specific genes were enriched for heat-shock proteins (*Hspa1a, Hspa1b, Hsph1, Hsp90aa1, Dnaja4*). While never described in this context, heat-shock proteins are known to play a role in response to hypoxia ^53^, an external cue that has been linked to T_RM_ development ^39^. Whether the up regulation of these genes in the LCMV model is related to hypoxic signaling or other external factors remains to be understood. In parallel, *Ikzf2* (encodes Helios) was identified as a skin-specific T_RM_ development transcription factor, which is best known for its ability to modulate regulatory T cell effector function and identity ^54,55^. Nonetheless, though *Ikzf2* was uniquely expressed in the skin dataset, we cannot rule out that this distinction was due to differences in the infectious agents used to induce T_RM_ phenotypes (VACV versus LCMV). Indeed, an *IKZF2*-expressing population of T_RM_ cells has been described in the human intestine ^56^. The T_RM_ cells isolated from the skin following VACV infection are those from the site of primary infection, where differentiation is driven in part by acute tissue inflammation. In contrast, the T_RM_ cells in the small intestine that arise from LCMV-infection delivered intraperitoneally represent those from a secondarily populated site, likely with much less inflammation. Such distinctions in the two infectious models that were analyzed could contribute to the differences in T_RM_-associated genes across tissue compartments.

Despite the differences in experimental design, tissues analyzed, and viruses used, 71 genes were commonly associated with T_RM_ development in both skin and siIEL. Of those 71 genes, 60 were defined as our “consensus” T_RM_ signature, after removing the 11 memory genes associated with T_CIRC_ populations. Previously reported T_RM_ signatures were derived from the comparison of T_RM_ to circulating memory T cells to define distinctions^10,20^. While these comparisons are valuable, our linear modeling approach enabled us to define T_RM_ differentiation at higher granularity by considering the effector T cell phase as our baseline. Additionally, our approach enabled us to compare the results of T_RM_ to that of T_CIRC_ to find genes common among different terminal memory T cell fates, regardless of circulatory capacity. Encouragingly, we did find overlap between our 60 gene consensus T_RM_ signature and those previously reported^10,20^. These included genes such as *Itgae*, *Nr4a3*, and *Rgs1*, all of which have been shown to play a role in T_RM_ formation^10,57^. It should be noted that our consensus signature lacks certain factors previously defined as T_RM_-specific genes, such as *Runx3, Notch1, Znf683* (encodes Hobit), and *Prdm1* (encodes Blimp1). *Runx3* was found to be a T_RM_-associated gene in the siIEL compartment, but not in skin. *Notch1* was found to have low expression in both the skin (8.7% of cells across all timepoints) and siIEL (11.7% of cells across all timepoints) datasets and its association with T_RM_ development has been best described in the lung^43^, suggesting it may be a tissue-specific driver of T_RM_ differentiation. *Prdm1* and *Zfp683* were both expressed at very low levels in our skin cells (*Prdm1* : 4% across all timepoints, *Zfp683* : 0.8% across all timepoints), making it difficult to evaluate their contribution in driving T_RM_ development through our linear modelling approach. However, it should be noted that *Prdm1* almost met our significance threshold for our siIEL T_RM_ gene list (regression slope = 0.199, p-value = 3.2e-07), supporting previous reports of this transcription factor’s involvement in T_RM_ development in the small intestine. Importantly, our 60-gene T_RM_ signature also includes genes uniquely captured by our linear modeling approach (e.g., *Fosb, Id3, Cebpb*) that could successfully distinguish resident from circulating T cell subsets, further validating our analytical strategy. Future work will be needed to characterize the role of these newly defined targets in T_RM_ development.

As noted above, this consensus T_RM_ signature included genes associated with responses to hypoxia as well as those classified as immediate early response genes. The immediate early genes were hallmarked by AP-1 transcription factor members, whose importance in T_RM_ differentiation across tissues was also highlighted in our SCENIC analysis of regulons. Additionally, motifs corresponding to AP-1 binding sites were found to be enriched in the open chromatin of T_RM_ when compared to T_CM_. Though the importance of AP-1 family members in T_RM_ development have been described recently in the siIEL^12^, our analyses confirmed that this is likely a generalizable marker of T_RM_ development and maintenance. This study is also the first to show AP-1-member protein nuclear staining in resting T_RM_ cells. One proposed hypothesis for the role of AP-1 transcription factors in T_RM_ differentiation is that it can suppress T-bet expression, a reported necessary step for T_RM_ development^19^. However, in both the skin and siIEL tissue niches, we saw a strong correlation between AP-1 family members and T-bet expression levels (encoded by the *Tbx21* gene). Given this observation, it remains unclear what role AP-1 plays in T_RM_ differentiation and maintenance. In non-adaptive immunity settings in murine epidermal stem cells, AP-1 members have been shown to be critical to orchestrate inflammatory memory, leaving keratinocytes poised for rapid recall responses^58^. There is good evidence that T cell receptor ligation by antigen in the context of MHC results in calcium influx and nuclear translocation of NFAT family members^59^. NFAT-AP-1 complexes are some of the most potent known super-enhancers of T cell cytokines and effector functions^59–63^. Given this, it is tempting to hypothesize that constitutive AP-1 expression and its nuclear localization enable T_RM_ cells to be “poised” for the rapid recall immune responses after TCR engagement alone, which results in NFAT nuclear translocation and formation of the NFAT/AP-1 transcription enhancer complex. If true, this is a novel mode of memory T cell activation.

## Methods

### Mice

Wild-type C57BL/6 mice were purchased from Jackson Laboratory. OT-I/Rag1^-/-^ /Thy1.1 mice were bred and maintained in the animal facility of Harvard Institute of Medicine, Harvard Medical School. All animal experiments protocols were approved by the Institutional Animal Care and Use Committee at Brigham and Women’s Hospital. All animal experiments were conducted in accordance with the guidelines from the Center for Animal Resources and Comparative Medicine at Harvard Medical School.

### Adoptive transfer and viral infection

For adoptive transfer, lymph nodes were collected from naïve OT-I/Rag1^-/-^ /Thy1.1 mice at the age of 6-8 weeks. OT-I T cells were then purified by negative magnetic cell sorting using mouse CD8α+ T-cell isolation kit (130-104-075; Miltenyi Biotec) according to the manufacture’s protocols. Purified OT-I cells were then transferred intravenously into gender-matched C57BL/6 recipient mice at the number of 5 × 10^5^ cells per mice. VACV-OVA was a kind gift from Dr Bernard Moss (National Institutes of Health, Bethesda, MD). VACV-OVA stocks were expanded in Hela cells (American Tissue Culture Company) and titrated in CV-1 cells (American Tissue Culture Company) by standard procedures. VACV-OVA were infected to mice at 2×10^6^ PFU/mice by skin scarification (s.s) on ear and tail as described previously^16,17^.

### Flow cytometry and cell sorting, and Imaging flow cytometry

Single cell suspensions were prepared as described before. Briefly, for lymph nodes, tissue specimens were mashed through 70µm cell strainers before lysed with RBC lysis buffer (00-4333-57; eBioscience). For skin, tail skin as well as separated dorsal and ventral halves of ear skin were minced and digested at 37 °C for 30 min, with HBSS solution with 1 mg/ml collagenase A (11088785103; Roche) and 40 μg/ml DNase I (10104159001; Roche) before being filtered through 70µm cell strainers. Cells were washed 3 times and kept in PBS supplemented with 2% FBS.

For flow cytometry, digested and purified skin single cells suspensions were stained and loaded onto FACSCanto II (BD Biosciences) for analysis or FACSAria (BD Biosciences) for sorting. To isolate OT-I cells for scRNAseq, cells were stained with antibodies against mouse CD8a (100714; Biolegend) and CD90.1 (202518, Biolegend). For the isolation of T_CM_ cells, cells were also stained with an antibody against CD62L (560514; BD). To analyze markers of our skin2 population, cells were additionally stained with antibodies against anti-mouse CD74 (561941, BD), anti-mouse MHC-II (11-5322-82, eBioscience), anti-mouse DAP12 (bs-12630R-PE, Bioss) and anti-mouse FcεRI γ subunit (06-727, Sigma). FACS data were analyzed with Flowjo software (Tree Star). For Imaging flow cytometry, purified single cell suspensions were first sorted for CD90.1+ CD8+ cells, for a total yield of 24,000 cells. The sorted cells were then fixed in 20µl fixation buffer and loaded onto the Amnis ImageStreamX imaging flow cytometer. Pictures were then taken for three populations of cells distinguished by their level of CD74 expression (CD74^neg^, CD74^int^ and CD74^high^).

### T-CyCIF imaging

FFPE sections of mouse tail skin were prepared and t-CyCIF was performed as previously reported^29,64^ following the published protocol on protocols.io (dx.doi.org/10.17504/protocols.io.bjiukkew). Slides were stained with Hoechst 33342 (0.25 μg/mL; LI-COR Biosciences) and antibodies against CD8a (83012BC; Cell Signaling Technologies), CD11c (64675BC; Cell Signaling Technologies), JunB (3753; Cell Signaling Technologies), and cFos (sc-166940 AF647; Santa Cruz Biotechnologies) in SuperBlockTM Blocking Buffer. Images were acquired using the CyteFinder II HT Instrument (RareCyte Inc. Seattle WA) with a 20x/0.75 NA objective. ASHLAR (Alignment by Simultaneous Harmonization of Layer/Adjacency Registration) software was used to stitch the image tiles and register each immunofluorescence cycle together into a single OME-TIFF file.

### Single cell RNA-sequencing (scRNAseq)

For the scRNAseq profiling, live CD8a+CD90.1+ cells were sorted as described above and then approximately 12,000 single cells were loaded to each 10X channel with a recovery goal of 6,000 single cells. Cell suspensions were loaded along with reverse transcriptase reagents, 3’ gel beads, and emulsification oil onto separate channels of a 10X Single Cell B Chip, which was loaded into the 10X Chromium instrument to generate emulsions. Emulsions were transferred to PCR strip tubes for immediate processing and reverse transcription. Library preparation was performed according to manufacturer’s recommendations. Expression libraries were generated using the Chromium Single Cell 3’V3 chemistry (10X Genomics PN-120262). DNA and library quality was evaluated using an Agilent 2100 Bioanalyzer and concentration was quantified using the Qubit dsDNA high-sensitivity reagents (ThermoFisher). Gene expression libraries were sequenced on an Illumina NextSeq instrument using the Illumina NextSeq 500/550 with the following sequencing configuration: Read 1=28, Read 2=56, index 1=8, index 2=0.

### Read alignment and quantification

Raw sequencing data was pre-processed with CellRanger (v3.0.2, 10X Genomics) to demultiplex FASTQ reads, align reads to the mouse reference genome (mm10), and count unique molecular identifiers (UMI) to produce a cell x gene count matrix ^65^. All count matrices were then aggregated with Pegasus (v0.17.2, Python) using the *aggregate matrices* function ^66^. Droplets with >20% mitochondrial UMI or <500 unique genes detected were deemed low-quality cells or empty droplets and were filtered out of the matrix prior to proceeding with downstream analyses. The counts for each remaining cell in the matrix were then log-normalized by computing the log1p (counts per 100,000), which we refer to in the text and figures as Log(CPM).

### Cell clustering

For all cell clustering analysis, 2,000 highly variable genes were selected using the *highly_variable_features* function in Pegasus and used as input for principal component analysis^66^. To account for technical variability between batches, the resulting principal component scores were aligned using the Harmony algorithm^67^. The top 50 principal components were used as input for generating a neighborhood graph. Clustering the neighborhood graph was performed using the Leiden algorithm^68^ and the data was represented using the Uniform Manifold Approximation and Projection (UMAP) algorithm (spread=1, min-dist=0.5)^69^ and force-directed layout embedding using the ForceAtlas2 algorithm^70^.

### Doublet and non-CD8 T cell removal

For all iterations of clustering analysis, cell clusters that were likely to represent cell doublets and/or non-CD8 T cell contaminating populations were filtered using a biologically informed approach. Clusters that had < 60% of cells expressing *Cd3d, Cd8a* and *Trbc2* were removed from the analysis. Additionally, we removed a cluster of cells isolated from lymph nodes that had high expression of both T cell markers (*Cd3d, Cd8a, Trbc2*) and B cell markers (*Ms4a1, Cd79a, Cd19*); these cells were marked as doublets and removed from downstream analyses.

### Marker gene identification

The marker genes defining each distinct cell cluster from our skin clustering analysis was determined by applying two complementary methods. First, we captured genes with high expression in each cluster by calculating the area under the receiver operating characteristic (AUROC) curve for the Log(CPM) values of each gene as a predictor of cluster membership using the *de_analysis* function in Pegasus. Genes with an AUROC ≥ 0.75 were considered marker genes for a particular cluster. Second, we created a pseudobulk count matrix to identify genes with lower expression that were highly specific for a given cluster^71^. Specifically, we summed the UMI counts across cells for each unique cluster/sample combination to create a matrix of *n genes x (n samples*n clusters)* and performed “one-versus-all” (OVA) differential expression (DE) analyses for each cluster using the *DESeq2* package (v1.32.0, R v4.1.0)^72^. For each cluster, we used an input model *gene ∼ in_clust*, where *in_clust* is a factor with two levels indicating if the sample was in or not in the cluster being tested. A Wald test was then used to calculate *P* values and compute a false discovery rate (FDR) using the Benjamini-Hochberg method. We identified marker genes that were significantly associated with a particular cluster as having an FDR<0.05. Marker genes (excluding ribosomal and mitochondrial genes) for each cluster were sequentially identified by first selecting genes with an AUROC ≥ 0.75, followed by those with an OVA pseudobulk FDR <0.05, and up to the top 30 genes were visualized with the *ComplexHeatmap* package (v2.8.0, R)^73^. The full list of marker genes is compiled in **Supplemental table 1**.

### Pairwise Spearman correlations between clusters

To compare the gene expression profiles of dLN and skin cell subsets, pairwise Spearman correlations between the OVA log2 fold-change values for all genes were calculated using the *cor* function in R. The resulting correlations were visualized with the *ComplexHeatmap* package (v.2.8.0, R).

### Waddington-OT trajectory analysis

Waddington-OT was run using the published optimal transport procedure as previously described within the CellRank (v1.5.1, python) suite of tools. Briefly, for a given set of cells *C* at timepoint *T*, “descendent” and “ancestor” distributions are estimated as the mass distribution of *C_T_* at later and earlier timepoints respectively. The mass distributions are estimated by transporting *C_T_* to the cells at the neighboring timepoints (*C_T + 1_, C_T-1_*), creating “temporal couplings”. The temporal couplings for all adjacent timepoints were calculated assuming a uniform growth rate. These temporal couplings were used to examine the relationship between different clusters across timepoints. Macrostates were calculated using the Generalized Perron Cluster Cluster Analysis (GPCCA) estimator with the *compute_macrostates* function in Cellrank. After defining our terminal states, absorption probabilities were calculated to estimate the probability of each cell reaching each terminal state using the *compute_absorption_probabilities* function in Cellrank. To define drivers of terminal differentiation, the absorption probabilities for each macrostate were correlated with gene expression using the *compute_lineage_drivers* function in Cellrank. The results of these correlations were reduced to only include transcription factors by referencing the mouse TFdb^74^.

### Linear modeling of gene expression

To define genes associated with T_RM_ development, cells from peripheral tissues (i.e., skin, siIEL) were used to create a pseudobulk count matrix where UMI counts were summed across cells for each unique timepoint to create a matrix of *n genes x n timepoints*. Linear modeling for each tissue was performed on the pseudobulk count matrix using the *DESeq2* package (v1.32.0, R v4.1.0). The input model was *gene ∼ timepoint* where *timepoint* was a continuous variable with integer values representing the order of timepoints (e.g., in skin : D2 = 0, D5 = 1, D10 = 3). Significant differentially expressed genes (DEG) were defined as those with an FDR < 0.1 (Wald test), were expressed in >5% of cells and had a regression slope > 0.2. Significant genes from these analyses were visualized with *ComplexHeatmap* package (v2.8.0, R). All linear model results can be found in **supplemental table 4**.

### SCENIC regulon analysis

To identify putative transcription factors associated with cell development, we employed the SCENIC workflow using Pyscenic (v0.11.2, Python)^44,45^. In short, this workflow first identifies potential transcription factor targets based on their co-expression. These modules of co-expressed genes then undergo cis-regulatory motif enrichment analysis in their corresponding promoters (within 500bp of the transcriptional start site) and are filtered to only include genes that have a corresponding motif. The mm9 motif database provided by the SCENIC authors were used for the motif enrichment analysis. A given transcription factor and filtered target genes are termed a “regulon”. The relative activity of these regulons was assessed using the AUCell methodology provided in the SCENIC workflow, where the enrichment of a regulon is calculated relative to non-regulon genes in each cell. To identify regulons associated with temporal development, the mean of the AUCell values for every regulon was taken for the cells from each timepoint to create an AUCell matrix of *n regulons x n timepoints*. We then employed linear modeling on the matrix using the *lm* function in R.

### Gene set scoring and Gene set enrichment analysis

Gene set scoring was performed using the *calc_signature_score* function from Pegasus. The “Immediate early gene” signature included the following 14 genes : *Fos, Fosb, Fosl2, Gem, Junb, Zfp36l1, Nr4a2, Nr4a1, Dennd4a, Ifrd1, Rel, Nr4a3, Egr1, Dusp1*. The “Hypoxia” signature was based on the HALLMARK hypoxia gene signature from MsigDB.

GSEA was performed using the *fgsea* function from the fgsea package (v1.18.0, R) with 10,000 permutations to test for independence. For GSEA performed to find gene sets associated with T_RM_ development, the input gene rankings were based on the linear modeling pseudobulk regression slope values, where the gene with the highest regression slope was ranked first and the lowest regression slope last.

### ATAC-seq analysis

Raw sequencing data was first aligned to the mouse reference genome with Bowtie2^75^. The resulting BAM files were then filtered to remove mitochondrial reads and PCR duplicates with Picard (v2.27.4, Java). Peak calling was performed for each sample with MACS2 using the BAMPE mode^76^. Motif analysis was performed using HOMER software (v4.11.1, Perl) with default settings^77^.

## Data and Material availability

scRNA-seq count matrices and related data will be available upon publication of this study.

## Code availability

Source code for data analysis will be available on GitHub upon publication of this study.

## Acknowledgements

This work was made possible by the generous support from the National Institute of Health (R01AR065807 and R01AI127654 to TSK; DP2CA247831 to ACV; U2C-CA233262 to PKS), and the Ludwig Cancer Research Center (to PKS) and the Massachusetts General Hospital Transformative Scholar in Medicine Award (to ACV).

## Author Contributions

TSK designed the overall study. YP, YY, JZ, TT, TP performed the mice experiments and sample collection. KM generated the scRNAseq data. YP and JZ helped generate the ATACseq data. Data analysis and interpretation was performed by NPS with substantial contributions from YY, YP, ACV, TSK. ACV managed and supervised scRNAseq data generation and analysis. YY performed follow-up validation experiments and immunophenotyping. JBW and SP performed the CyCIF experiments with guidance from PKS. NPS, YY, ACV, TSK wrote the manuscript with substantial revisions by JBW. All authors read or provided comments on the manuscript.

---

**Extended data figure 1.**
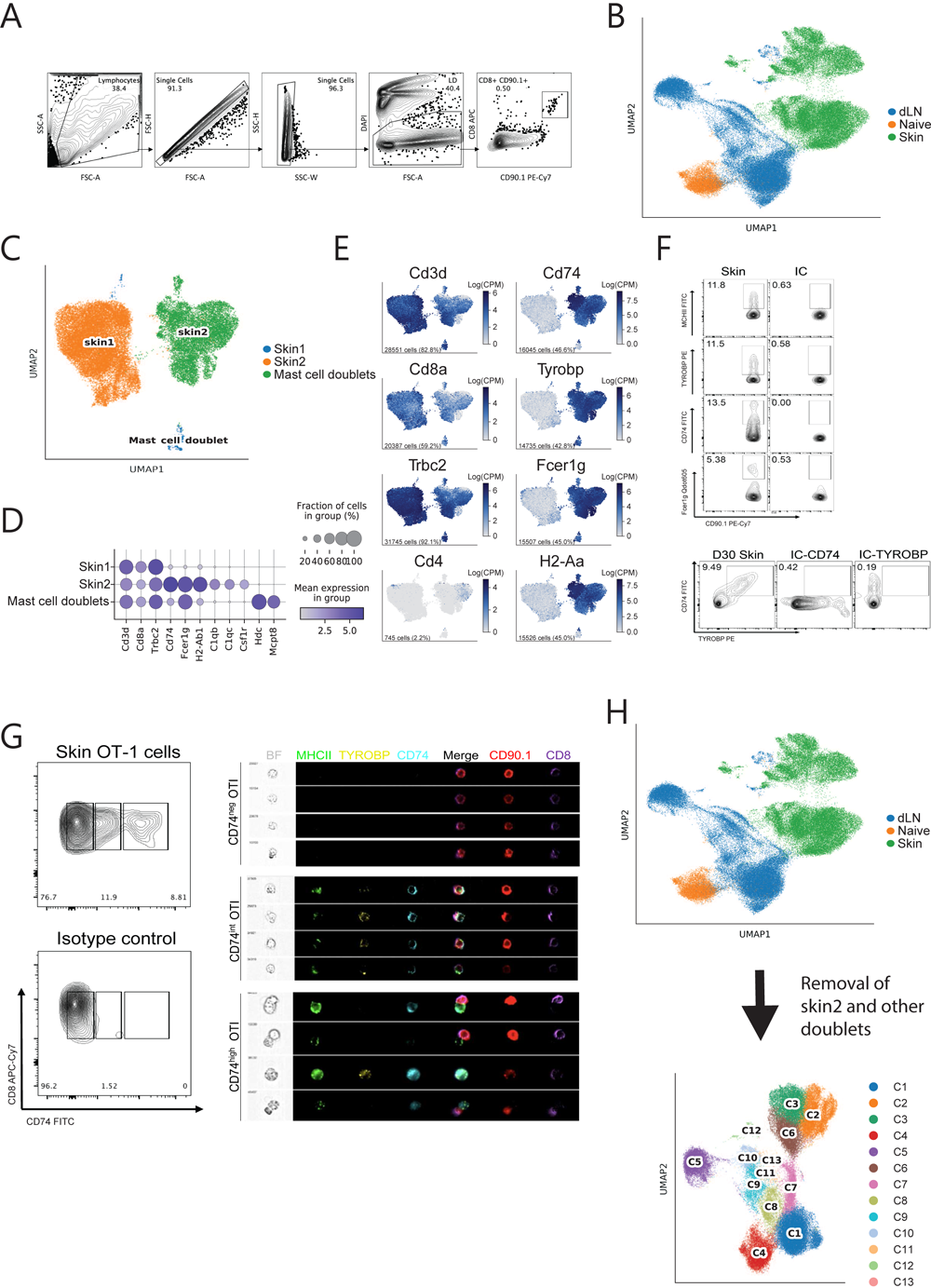
**(A)** Gating strategy used to isolate live CD8^+^CD90.1^+^ cells for downstream scRNAseq profiling. **(B)** Uniform Manifold Approximation and Projection (UMAP) embedding of 76,028 high-quality cells colored by tissue source. **(C)** UMAP embedding of 34,465 high-quality cells derived from skin, colored by major clusters segregating skin1 from skin2. **(D)** Dot plot showing the percentage (size of the dot) and scaled expression (color) of genes associated with each major cluster in **(C)**. **(E)** Feature plots using color to indicate gene expression levels (Log(CPM)) of canonical T cell genes as well as marker genes for the skin2 cluster. Cell number and percentage represent expression across all cells. **(F)** (top) representative flow cytometry analysis of D30 OT-I cells stained for anti-CD74, anti-TYROBP, anti-MHC-II, anti-FCER1G or isotype controls (Y-axis) and anti-CD90.1 (X-axis) to validate the existence of “skin2” cell subset. (bottom) Representative flow cytometry plot of D30 OT-I cells stained for anti-CD74 (Y-axis) and anti-TYROBP (X-axis) or isotype controls. **(G)** (left) Representative flow cytometry analysis of D30 OT-I cells isolated from skin stained for anti-CD8 (y-axis) and anti-CD74 or an isotype control (x-axis). CD8+ cells were gated into 3 distinct populations: CD74^neg^, CD74^int^ and CD74^high^. (right) Single-cell images of D30 OT-I cells grouped by their CD74 status (CD74^neg^, CD74^int^, CD74^high^) co-stained for MCH-II, TYROBP, CD74, CD90.1 and CD8 (ImageStream). **(H)** Schematic of our cell filtering process to remove suspected doublet and non-CD8 T cell contaminants. Given our inability to separate doublets from singlets in the skin2 cluster, all skin2 cells (along with other predicted doublet populations) were filtered from the scRNAseq dataset used for downstream analysis.

**Extended data figure 2.**
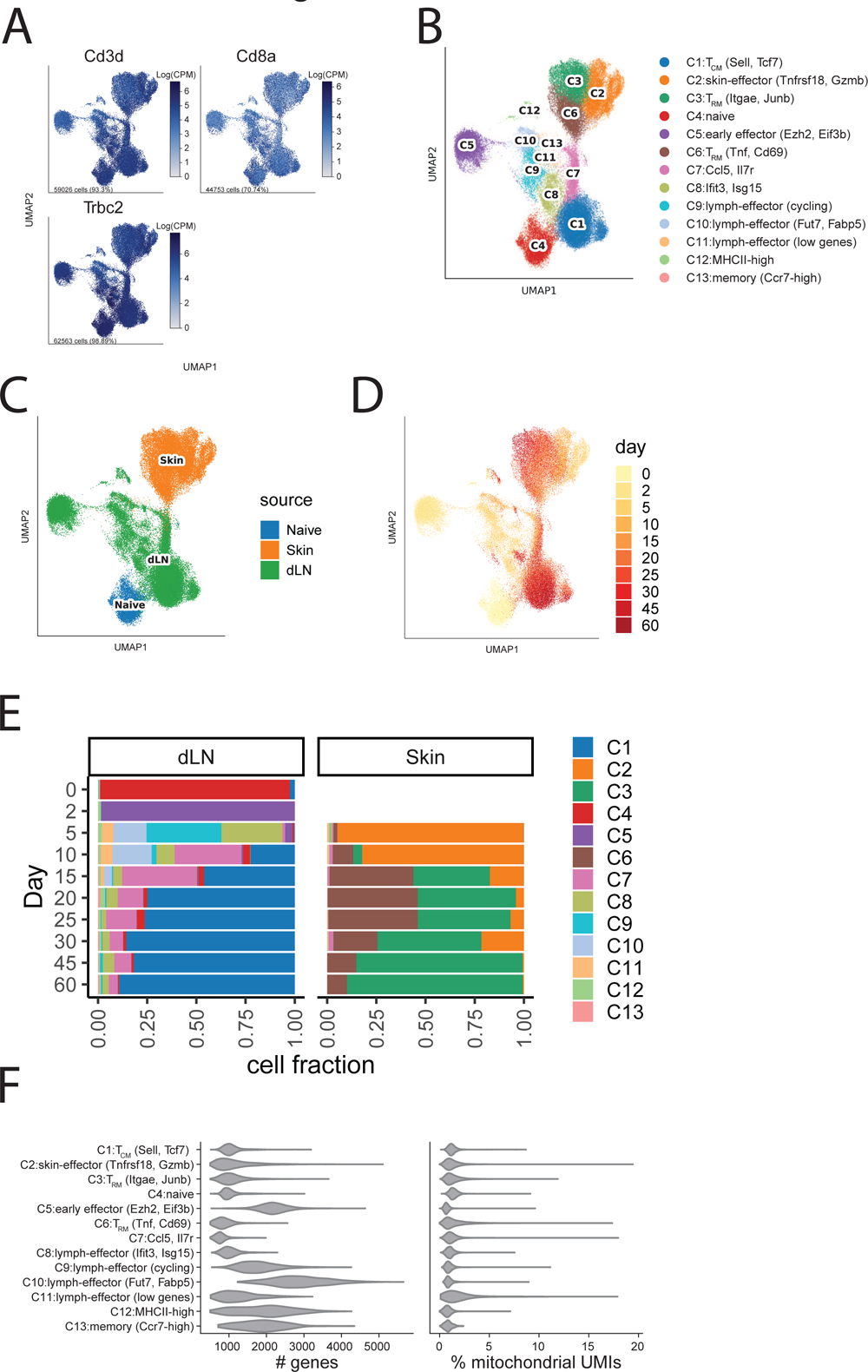
**(A)** UMAP embedding feature plots of 63,265 high-quality single cells, using color to represent gene expression levels (Log(CPM)) of canonical T cell genes (*Cd3d, Cd8a, Trbc2*). Cell number and percentage represent expression across all cells. **(B)** UMAP embedding showing predicted Leiden clusters listed on the right. **(C)** UMAP embedding of cells pseudocolored by tissue source. **(D)** UMAP embedding of cells pseudocolored by experimental timepoint. **(E)** Cluster composition at every timepoint. Bars represent the fraction of cells in dLN (left) and skin (right) that were assigned to the corresponding clusters. **(F)** Distribution of the number of captured genes (left) and percentage of mitochondrial unique molecular identifiers (UMIs) (right) in the cells across all clusters.

**Extended data figure 3.**
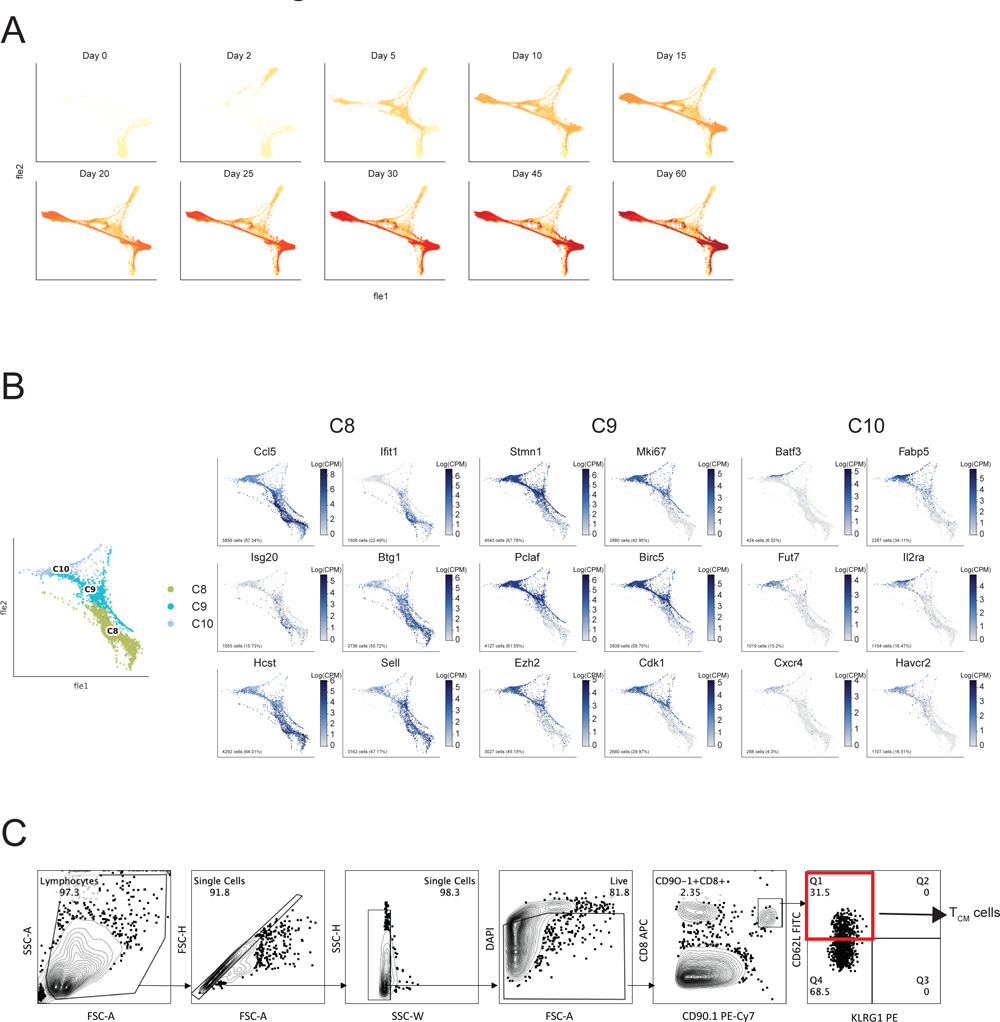
**(A)** Force-directed layout embedding (FLE) of single-cells colored by timepoint. Each plot in the panel represents the cells from that timepoint and all previous timepoints. **(B)** (left) FLE of clusters most associated with day 5 dLN cells (C8, C9, and C10). (right) Feature plots using color to represent gene expression levels (Log(CPM)) of genes associated with each cluster. Cell number and percentage represent expression across all cells in C8, C9 and C10. **(C)** Gating strategy for the isolation of T_CM_ cells used for ATACseq profiling experiment. Gated T_CM_ cells are highlighted in the red box.

**Extended data figure 4.**
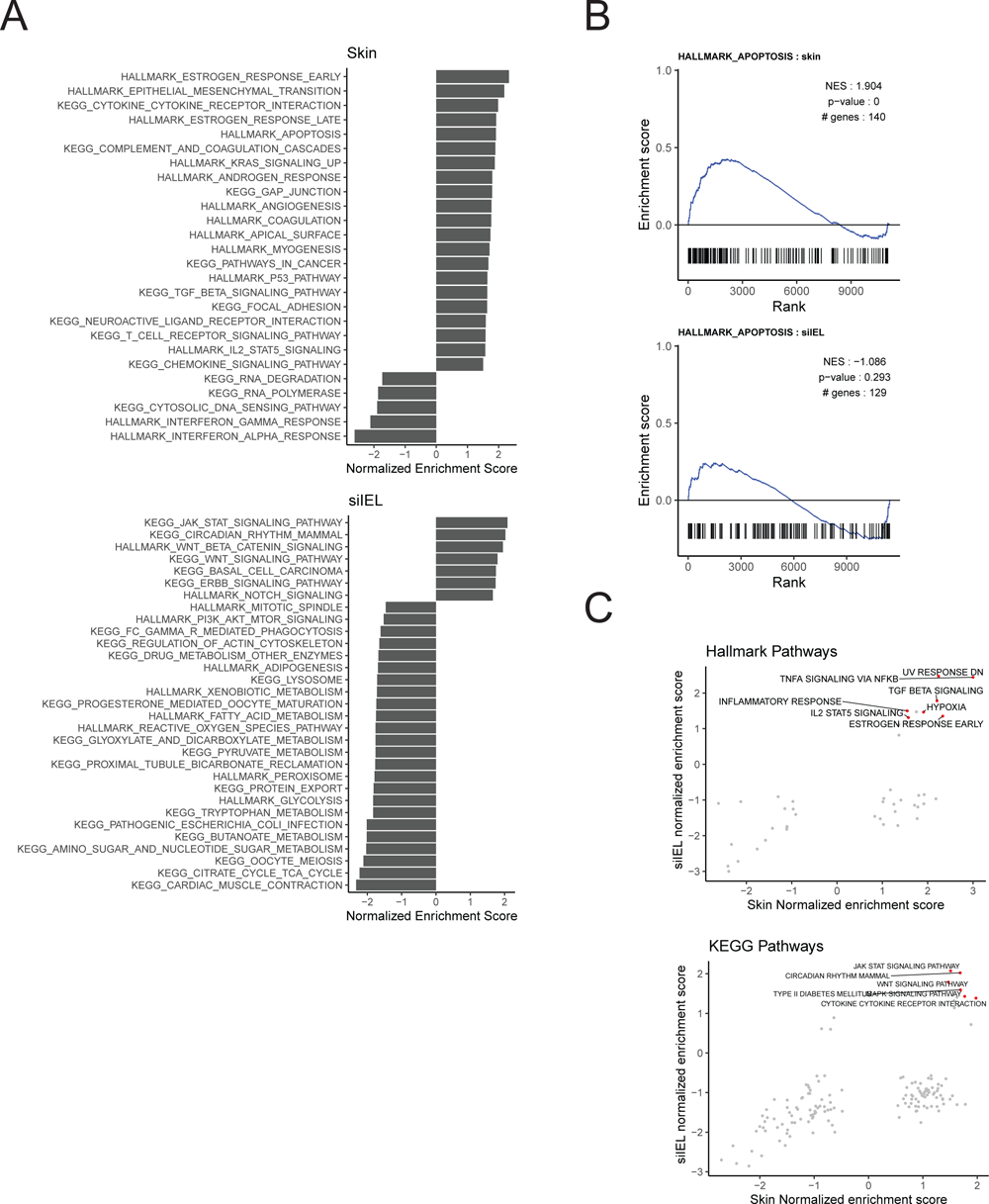
**(A)** Gene set enrichment analysis (GSEA) of both skin and siIEL T_RM_ genes was performed using the KEGG and HALLMARK gene set libraries. Plots show the normalized enrichment scores (x-axis) of all gene sets that were uniquely significant in skin (top) and siIEL (bottom) (FDR < 0.1). **(B)** GSEA of the HALLMARK apoptosis gene signature for skin (top) and siIEL (bottom) T_RM_ genes. **(C)** Gene set enrichment analysis results for skin and siIEL using the HALLMARK (top) and KEGG (bottom) pathways. Each point represents a gene set and the x-and y-axes represent the skin and siIEL normalized enrichment scores respectively. Labelled points highlighted in red show gene sets that were significantly associated with T_RM_ gene lists in both skin and siIEL (FDR < 0.1).

**Extended data figure 5.**
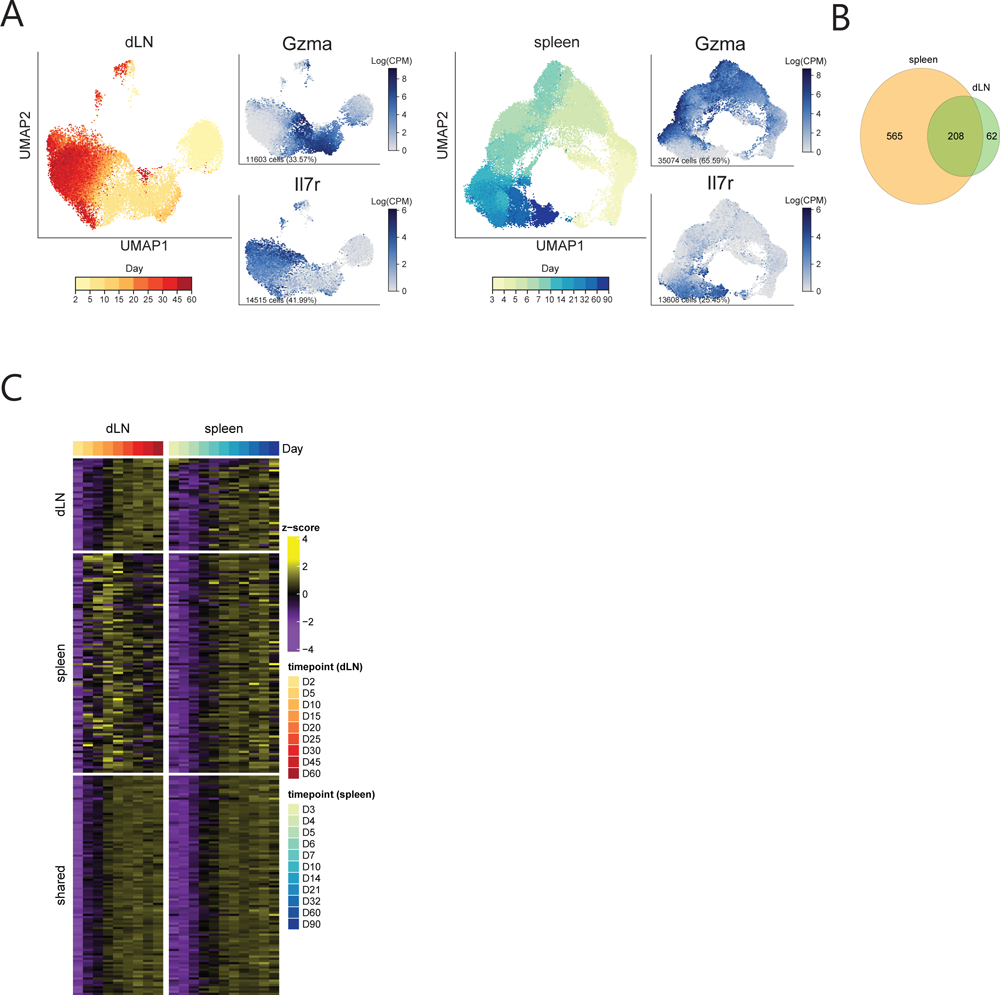
**(A)** UMAP embedding of dLN (left) and publicly available spleen data (right) pseudocolored by experimental timepoint. To the right of each timepoint UMAP are feature plots using color to indicate gene expression levels (Log(CPM)) of *Gzma* and *Il7r*. Cell number and percentage represent expression across cells of all timepoints within the given anatomical niche. **(B)** Venn diagram of the significant T_CIRC_ -associated genes in spleen (left) and dLN (right), as determined by linear modeling. **(C)** Heatmap showing the top genes unique to the dLN (top, n = 62 genes) and spleen (bottom, n = 100 genes) T_CIRC_ signatures. Top bar indicates the associated timepoints. Color scales denote the normalized gene expression (mean zero, unit variance) for each timepoint.

**Extended data figure 6.**
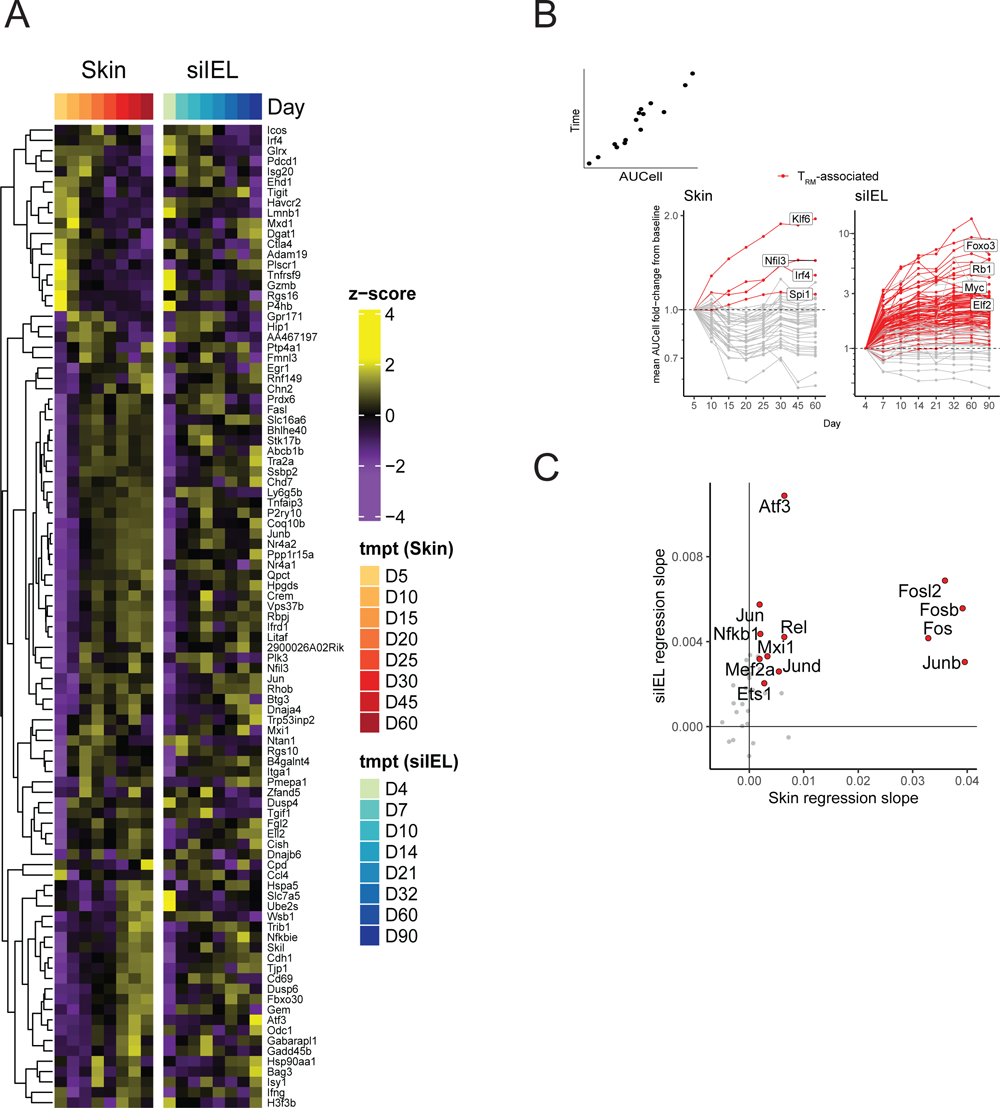
**(A)** Heatmap showing the expression of genes unique to the Milner *et al.* T_RM_ gene signature across timepoints in the skin (left) and siIEL (right) datasets. Color scale denotes normalized gene expression (mean zero, unit variance) for each timepoint. **(B)** SCENIC regulon analysis of skin (left) and siIEL (right) cells. Each line represents a regulon. The x-axis represents the timepoint and the y-axis represents the fold-change of the mean AUCell value of the given regulon at that day when compared to the baseline timepoint (skin = D5, siIEL = D4). Red lines denote significant regulons unique to either skin or siIEL T_RM_ development. **(C)** SCENIC regulon analysis of skin and siIEL cells. Each point represents a regulon and the x-and y-axes represent the regression slope of the mean AUCell score of the regulon in skin and siIEL over time respectively (see **Methods**). Red points denote regulons that were significantly associated with both skin and siIEL T_RM_ development (FDR < 0.1, linear model).

**Extended data figure 7.**
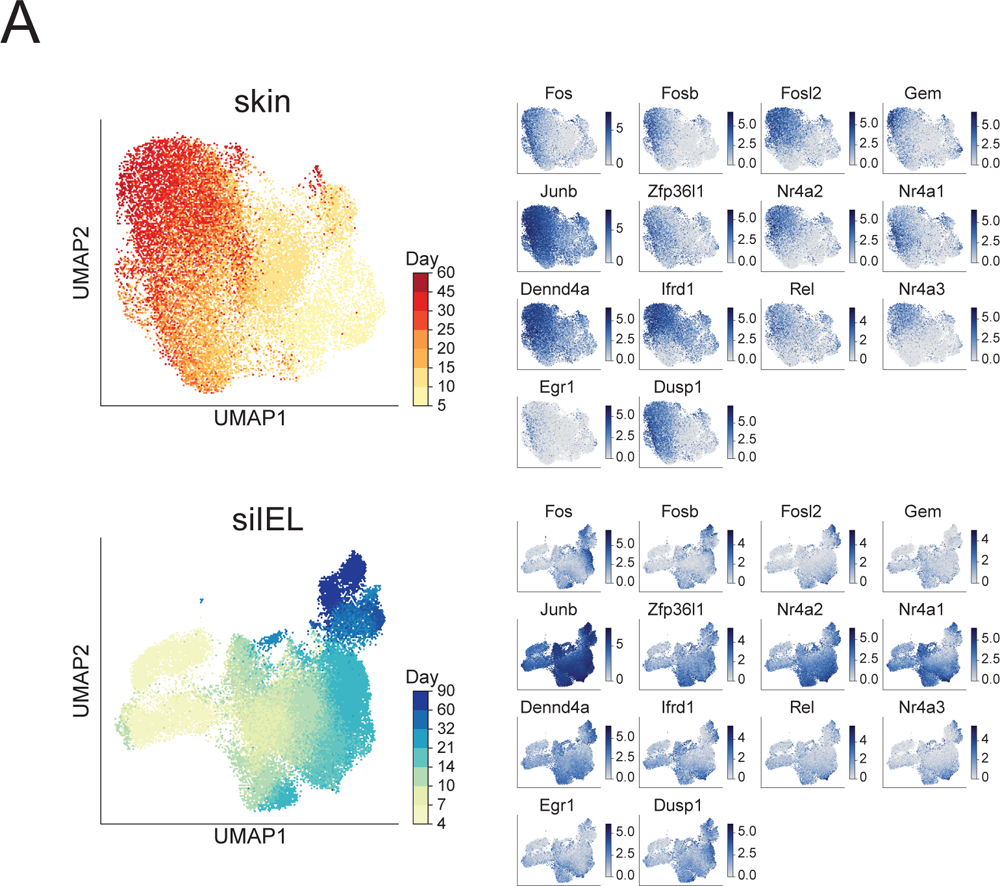
**(A)** (left) UMAP embedding of skin (top) and publicly available siIEL scRNAseq data (bottom) pseudocolored by experimental timepoint. (right) Feature plots using color to represent gene expression levels (Log(CPM)) of known immediate early genes.

